# Predicting improved protein conformations with a temporal deep recurrent neural network

**DOI:** 10.1101/275008

**Authors:** Erik Pfeiffenberger, Paul A. Bates

## Abstract

Accurate protein structure prediction from amino acid sequence is still an unsolved problem. The most reliable methods centre on template based modelling. However, the accuracy of these models entirely depends on the availability of experimentally resolved homologous template structures. In order to generate more accurate models, extensive physics based molecular dynamics (MD) refinement simulations are performed to sample many different conformations to find improved conformational states. In this study, we propose a deep recurrent network model, called DeepTrajectory, that is able to identify these improved conformational states, with high precision, from a variety of different MD based sampling protocols. The proposed model learns the temporal patterns of features computed from the MD trajectory data in order to classify whether each recorded simulation snapshot is an improved conformational state, decreased conformational state or a none perceivable change in state with respect to the starting conformation. The model is trained and tested on 904 trajectories from 42 different protein systems with a cumulative number of more than 1.7 million snapshots. We show that our model outperforms other state of the art machine-learning algorithms that do not consider temporal dependencies. To our knowledge, DeepTrajectory is the first implementation of a time-dependent deep-learning protocol that is re-trainable and able to adapt to any new MD based sampling procedure, thereby demonstrating how a neural network can be used to learn the latter part of the protein folding funnel.

## Introduction

Protein structure prediction from sequence tries to overcome the limitations of experimental structure determination, which are often time consuming and infeasible for certain types of proteins. Furthermore, construction of protein models seems to be the only practical solution for structural genomics where a high rate of newly discovered protein sequences demands for automated structure determination [1]. Current stateof-the-art methods which make use of template based modelling (TBM) are partially successful [2–8]. However, the quality of these TBMs is completely dependent on the presence of homologous proteins where the structure has been experimentally determined. An extension to TBM are so-called physics-based refinement methods that further try to improve the initial models by extensively sampling new conformations; essentially, emulating the later part of the protein folding pathway. Methods which make use of conformational sampling with molecular dynamics (MD) simulations with an allatom physical force field have proven to be successful in sampling improved conformational states. Currently, the most successful refinement simulations are based on multiple replicated simulations in the nanosecond scale with position restraints on parts of the protein to prevent drifts [9–11]. Yet, the most challenging problem is the reliable identification of improved quality configurations from this time-series trajectory data, from millions of possible solutions [12–14].

The continued progress in deep-learning research has demonstrated success for a number of noisy sequence or time-series problems [15–17]. In this work, a temporal deep-learning model for snapshot classification of MD trajectory data is formalized that makes explicit use of the time-dependent nature of MD based trajectory data. In particular, the interest lies in whether it is possible to identify when, or if, improved quality conformations of a protein are reached, from a variety of starting model qualities. Progress in this area is important for high accuracy model building that is, for example, required for biomolecular understanding of protein function and *in-silico* rational drug design. From the generated trajectory of conformational snapshots, predictions about a protein’s conformational states are based on energies and distance metrics in time. To this end, a deep recurrent neural network (RNN) [18] with gated recurrent units (GRUs) [19] is trained to classify each snapshot into one of three classes: improved quality, no-change in quality, and decreased quality. The change of quality is defined as an increase or decrease in the global distance test total score (GDTTS) [20,21] from the starting configuration, as measured with respect to the reference crystal structure.

The results show that it is possible to train a RNN model that identifies improved and decreased quality states in different MD based refinement protocols with nanosecond time-scale. Furthermore, the proposed model outperforms classic machine learning models and deep learning models that do not consider temporal dependencies during their training task. To be precise, our model achieves a mean cross-validation precision on the improved state assignment of 41.5% compared to 14.0%, 12.1% and 0.0% for random forest (RF) [22], k-nearest neighbours (KNN) [23], and logistic regression (LR) [24], respectively. The results also show that a deep representation and temporal patterns learned by the RNN are important and contribute to a higher precision of identifying improved quality snapshots.

## Results

### The DeepTrajectory model

The DeepTrajectory model is summarized in Fig. 1. The model takes as input the computed features from the sampled MD trajectory data and tries to predict the state change relative to the starting configuration for each snapshot. The three classes learned by the classifier are improved state (*I*), no-change state (*N*) and decreased state (*D*), and reflect whether a more accurate conformation of the protein is reached with respect to the reference crystal structure (See Fig. 1A). The trajectory is quantified via 19 metrics. These are 17 different potential energy terms and scoring functions (quantifying the energetics of the protein structure); and 2 distance-metrics known as root mean square deviation (RMSD) and GDTTS (measuring the deviation to the starting configuration at time-step 0). Details about the features are summarized in SI Table S1. Essentially, the DeepTrajectory model learns the temporal patterns of the input features in order to distinguish correct fold state changes from incorrect fold state changes. The trajectory data from the different protein systems and sampling runs used for training is presented to the network in mini-batches of 60 ps (30 steps) that continue after each iteration of training until the end of the training data is reached. The DeepTrajectory model is an RNN with GRUs (Fig. 1B). Each 2 ps time-step is represented by a standardized feature vector containing the values of the 19 features. The predicted state assignment for every snapshot by the RNN, i.e. *I, N* and *D*, is expressed as a probability distribution and the class with the highest probability is used as the final prediction.

**Fig. 1:**
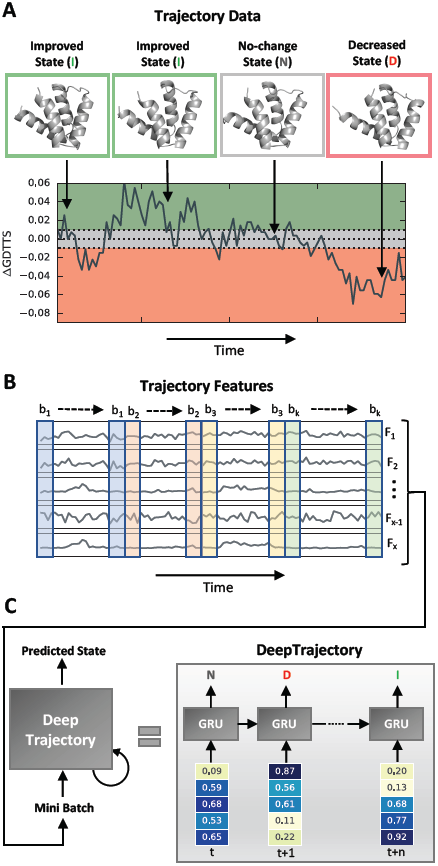
DeepTrajectory method overview. The method predicts improved conformational states, that more closely resemble the crystal structure observed conformation, in molecular dynamics (MD) trajectory data of template based models. (A) During sampling of the initially folded proteins with different MD based sampling protocols, transitions to conformations can be observed that represent an improved conformational state (ΔGDTTS *≥* 0.01), decreased conformational state (ΔGDTTS *≤ -* 0.01) and no-change in conformational state (|ΔGDTTS*|<* 0.01). Every 2 ps a snapshot of this trajectory is saved. (B) The trajectory of snapshots of different sampled conformational states is quantified by 19 features (*F*_1_,…, *F*_*x*_) that measure different energetic contributions or distance metrics of the protein structure at a particular time-point. (C) These temporal features are used to perform supervised mini-batch training (*b*_1_,…, *b*_*k*_) to train an RNN with several layers of GRUs that is able to classify each snapshot of the trajectory into three classes: improved state (*I*), no-change state (*N*), decreased state (*D*). Predictions for the trajectory of a new protein system, for which this state change assignment is unknown, are assigned by applying the trained DeepTrajectory model to every snapshot. Full details of the model and training procedure are available in the SI text.

### Data-set and performance criteria

The performance of DeepTrajectory was compared to a RF classifier, a KNN classifier and a LR classifier. The data used for training and testing accumulates to 904 trajectories and a total simulation time of 3419 ns from 42 different protein monomers. The used protein systems and their starting model quality were collected from the refinement category in rounds 11 and 12 from the Critical Assessment of protein Structure Prediction (CASP) experiment. The dataset consists of a wide range of GDTTS values from 0.3 to 0.9 (Fig. 2A). The sampled ΔGDTTS, that expresses the relative change in model quality relative to the starting model, ranges from *-*0.3 to 0.12 (Fig. 2B), details for each trajectory are shown in SI Table S5. The class assignment of each snapshot, shown in Fig. 2C, into one of the three different states *I, N* and *D*, have a relative distribution of 8.2, 14.5 and 77.3 percent, respectively. An analysis of the trajectory data as a Markov chain model shows the transition probabilities between the different states (Fig. 2D). These show that increased and decreased conformational states are more stable with a probability of 0.806 and 0.947 to remain in the same state, compared to no-change state with 0.626. This is also expressed by the observation that these states sample with higher frequency longer continuous segments, see SI Fig. S3A (improved state) and SI Fig. S3C (decreased state).

**Fig. 2:**
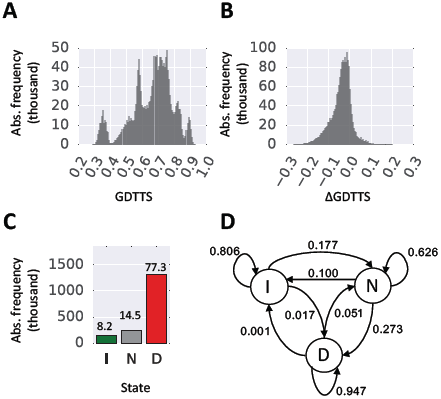
Trajectory dataset. (A) Histogram of GDTTS in the trajectory data. (B) Histogram of ΔGDTTS in the trajectory data. A positive value indicates an improvement and a negative value a decrease in model quality. (C) Absolute frequency of the three different states improved, *I*, no-change, *N*, and decreased, *D*. (D) Markov chain model of the three states improved, *I*, no-change, *N*, and decreased, *D*, visualised as circles and their directed transition probabilities shown as labelled arrows.

The aim of the classifier is to learn a temporal model in order to identify improved conformational states (*I*) with high precision, and decreased conformational states (*D*) with high recall. Thus, during training we are trying to minimize the number of false positive predictions for the improved state and the number of false negative predictions for the decreased state class. This results in a model with higher confidence that the predicted *I* state is correct and that a large number of *D* states could be identified. Additionally, the metric F1 is computed that is the harmonic mean of precision and recall. We compare this to all three base-line machine learning models. A detailed definition of the used performance metrics is given in the SI text.

### Comparison of model performance to other classifiers

The bar-plots seen in panels Fig.3A-C quantify the classification performance for all three classes *I, N* and *D* of the temporal RNN model and compare it to KNN, RF and LR. Most notably is the prediction performance of the RNN for the improved state. The RNN is able to identify improvements in folds with a markedly better precision than classical machine learning models. To be precise, the mean cross-validation precision for RNN, KNN, RF and LR have values of 0.415, 0.121, 0.139 and 0.000, respectively. For recall on the improved class the values are 0.037, 0.065, 0.001 and 0.000 for RNN, KNN, RF and LR, respectively. Values for the decreased class performance are similar for all models (Fig. 3C). For this class the models RNN, KNN, RF and LR produce a mean cross-validation precision of 0.790, 0.798, 0.778 and 0.777, respectively. The recall is 0.960, 0.888, 0.985 and 0.987 for RNN, KNN and RF and LR, respectively.

**Fig. 3:**
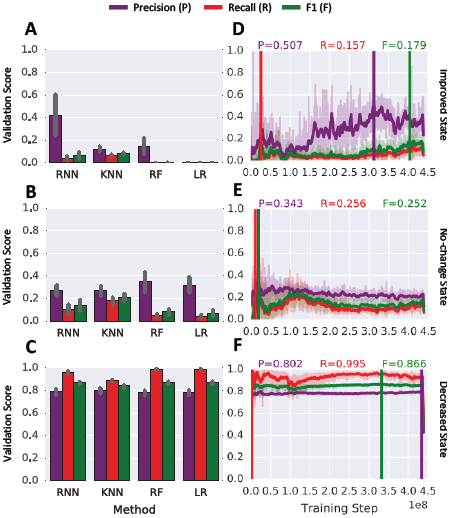
RNN performance comparison. (A)-(C) Comparison of validation-set performance for the 3 different states improved, no-change and decreased. Shown are the recurrent neural network (RNN) against k-nearest neighbours (KNN), random forest (RF) and logistic regression (LR) on the full cross-validation sets. The shown performance is the mean value of all seven validation sets. The error bars indicate the standard deviation computed from all 7 folds. (D)-(E) Classification precision, recall and F1 as a function of training steps for the three different states. Shown is the training progress for CV fold 4. The vertical lines indicate the best performance for each metric based on a moving average with a window-size of 30.

The confusion matrix (CM) in SI Table S3 shows the miss-assignment of predicted classes versus the actual class for all 4 tested models for validation-fold 4 of the cross validation (CV). For example, the RNN model predicted the correct true positive assignment for improved snapshots 1636 times and assigned the label improved incorrectly 314 times to no-change snapshots and 1653 times to decreased snapshots (see SI Table S3A). Compared to the three other models KNN, RF and LR this represents a notably better performance at identifying improved snapshots. For KNN, seen in SI Table S3B, a similar number, i.e. 1556, of true positive improved snapshot assignments compared to RNN could be achieved. However, this comes with a large number of false positive assignments where the KNN incorrectly assigns the improved label to 2386 nochange snapshots and 6690 decreased snapshots. The RF model is hardly predicting the improved class (SI Table S3C). This model generates 34 true positive assignments for the improved class. The number of false positive predictions, where the actual class is different from the predicted, is 25 and 39 for no-change and decreased, respectively. The LR model is not able to predict improved snapshots at all, i.e. the number of assignments is zero (SI Table S3D).

Fig. 3D-F shows the validation score for precision, recall and F1 as functions of training steps for the three classes (*I, N* and *D*). The improved class shown in Fig. 3D indicates that several million training steps are necessary to reach the best running average precision of 0.507. The best precision and recall for classes nochange (Fig. 3E) and decreased (Fig. 3F) are reached early on during training. Furthermore, for these two classes the validation score stays stable during the 300 epoch training process.

### Analysis of prediction performance

Fig. 4A shows DeepTrajectory’s true positive (TP), false negative (FN) and false positive (FP) predictions as cumulative distributions of ΔGDTTS for the improved state class. The results show a marked increase of FP predictions as ΔGDTTS tends towards the lower bound of the increased state definition (ΔGDTTS = 0.01). TP predictions have the highest increase in density for ΔGDTTS in the range from 0.01 to 0.05. For FN predictions of the improved state, the curve is less steep in the range 0.025 to 0.1 compared to TP, indicating fewer FN assignments. The no-change state cumulative distribution of FP predictions shows a rapid increase in the ΔGDTTS range from −0.05 to −0.01, with only a shallow increase for 0.01 to 0.05 (SI Fig. S4A). The cumulative distribution of FN and TP follow the same trend for no-change state predictions (SI Fig. S4A). For decreased state predictions most FP are distributed in the range from ΔGDTTS −0.01 to 0.01. The cumulative distribution of FN stays below the distribution of TP for the ΔGDTTS range −0.15 to −0.01 (SI Fig. S5A).

**Fig. 4:**
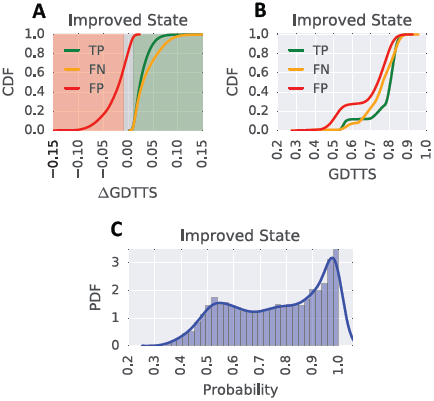
Performance analysis of DeepTrajectory. (A) Cumulative distribution of true positive (TP), false negative (FN) and false positive (FP) predictions for the improved state as a function of ΔGDTTS. The background colours red, gray and green indicate the ΔGDTTS regions for the improved, no-change and decreased states, respectively. (B) Cumulative distribution of true positive (TP), false negative (FN) and false positive (FP) predictions for the improved state as a function of GDTTS. (C) Shows the distribution of assigned probabilities for improved state predictions.

The cumulative density of TP, FN and FP as a function of absolute GDTTS is visualized in Fig. 4B for improved state predictions. A more rapid increase in FP predictions for GDTTS in the range 0.45 to 0.7 as compared to TP and FN is observed. This indicates that correct predictions of lower starting model qualities are less likely. However, TP predictions markedly increase for GDTTS in the range 0.7 to 0.85, showing the successful prediction of the descent to the native state of the latter part of the folding funnel. The cumulative TP distribution of the no-change state shows a similar behavior, where only higher quality models with GDTTS of 0.5 or more produce an increase in TP (SI Fig. S4B), whereas the decreased state distribution of TP, FN and FP shows a linear growth for all three curves (SI Fig. S5B).

In order to obtain an understanding of the confidence of DeepTrajectory to assign the three different states the predicted probabilities are considered as a variable and the probability density function of these probabilities were constructed from the whole data-set. Our model produces a wide range of assigned probabilities for predicted classes improved and no-change state (Fig. 4C and SI Fig. S4C) where the majority of probabilities are uniformly distributed from values of 0.5 to 0.9, with a slight increase in density in the range from 0.9 to 1.0. The probability density function for predicted decreased states is markedly different, most of the RNN’s computed probabilities are observed in the range from 0.9 to 1.0, whereas the range from 0.4 to 0.9 has a low density (SI Fig. S5C).

### Temporal and deep representation is important for precision and recall

We analyzed how important the temporal aspect of our RNN for classification success is. An RNN with sequence length of 1 (Fig. 5A) and 5 (Fig. 5B) experience training instability and a drastic drop in precision and recall for the improved state class compared to a model of length 30 (Fig. 5C) and 50 (Fig. 5D). The depth of the RNN is an important aspect too. An RNN with 1 (Fig. 5E) or 2 (Fig. 5F) layers results in a drop in precision and recall compared to 3 layers (Fig. 5G). Furthermore, a drop is observed when 10 layers (Fig. 5H) are used within the tested training time of 300 epochs.

**Fig. 5:**
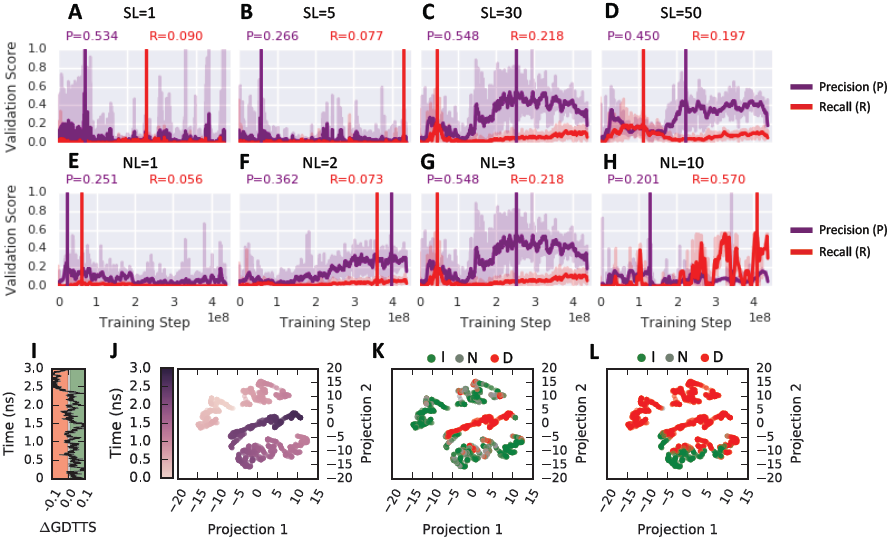
(A)-(D) Effect of sequence lengths 1, 5, 30 and 50 on precision and recall for the improved state class as a function of training steps tested on validation fold 4. (E)-(H) Effect of number of hidden layers with values 1, 2, 3 and 10 on precision and recall for the improved state class as a function of training steps tested on the 4th validation fold. The vertical lines indicate the best performance based on a moving average with a window-size of 30. (I) Trajectory trace of target TR821, the ΔGDTTS is shown as a function of its 3ns simulation time. Sampling of the green colored regions indicate an improved state, grey regions a no-change state and red a decreased state. Visualizations of the last hidden layer where the data-points are colored by (J) time, ground truth and (L) predicted labels. The 1024-dimensional last hidden layer was projected to 2-dimensions with t-SNE.

In order to better understand how the temporal aspect of our trajectory data is learned by the RNN, we analysed the output of the last hidden layer. We examined the internal feature representation learned by the RNN using t-distributed Stochastic Neighbour Embedding (t-SNE) [25]. As an example, a trajectory of one protein system is visualised. In Fig. 5I, the ΔGDTTS change as a function of time for a 3ns simulation is shown. The learned representation by the RNN of this trajectory is shown in Fig. 5J-L, where each point represents a projection from 1024 to 2 dimensions from the output of the last hidden layer. The projection in Fig. 5J is coloured by simulation time. We can see that points close in time also cluster together close in the projected space. The projections in Fig. 5K and L are coloured by the ground truth label and the predicted label, respectively. The projections of the predicted improved class shows pattern of aggregation into the same region.

## Discussion

The results show that the proposed RNN model with GRU cells is able to outperform classical machine learning methods such as RF, LR and KNN, which are representative of state-of-the-art classical machine learning algorithms and have been successfully applied to other bioinformatic domains [26–28]. In particular, the model presented here achieves a mean precision of 0.415 on the validation set of the CV compared to 0.121, 0.140 and 0.000 for KNN, RF and LR, respectively. This suggests that learned temporal dependencies of the used energy terms and distance metrics as input features are important to identify sections of fold improvements in the trajectory. This claim is further supported by inspection of the transition probabilities between *I, N* and *D* in Fig. 2D that are not random. The transition probabilities of staying in their respective states from time-step t to t+1 are 0.806, 0.626 and 0.947 for *I, N, D* respectively, and indicates that sequential state awareness is important for this particular classification problem.

A major application of DeepTrajectory would be in on-line classification of MD refinement simulations. Here, the model would be applied in parallel to a MD simulation to classify each snapshot of the trajectory. This would allow to probe whether the sampling has yielded enough improved conformational states and can be stopped, or vice versa, if no snapshots are classified as improved during the sampling run the simulation could be continued or restarted with different conditions. Moreover, the presented methodology could be extended to complete folding simulations by guiding the sampling process from a partially folded state to a near native state [29]. For example, this could be used in combination with molecular dynamics and Markov-state models in order to classify micro-states [30 31]. This would allow for selecting the correct micro-states that point to a descent of the folding funnel. Finally, similar to the RNN applied to trajectories of protein monomers, the proposed model could be trained on the trajectories of protein-protein assemblies [32 33]. Input features for such a method would need to focus on molecular descriptors that specifically describe protein-protein interactions [34 35].

We believe DeepTrajectory makes an important contribution to the field of protein structure refinement, an important area in structural bioinformatics with applications in rational drug design [36] and systems biology [37]. Our work has shown that a temporal model based on a RNN can predict, with high precision, the state changes to improved protein conformations. This opens the door for more application driven research exploiting deep learning as means to improve accuracy of protein structure prediction.

### Materials and Methods

The DeepTrajectory model is described in Fig. 1. The model is implemented and tested on the TensorFlow Python library in version 1.0.0 [38]. A detailed description of the model is available in the SI text. The used 904 trajectories originate from MD simulations of 42 different protein systems; detailed sampling protocols are described in the SI text.

The training and testing of the DeepTrajectory model is based on a seven fold cross-validation where in each fold the trajectories for 6 protein systems are selected for validation and 36 for training. In total a cumulative number of *≈* 1.7 million snapshots are used. Training of the model is done in mini-batches that are continued without overlap in each training step until the epoch is finished. During training, the model is provided with a feature vector of size 19, that consists of 17 different energy functions, or terms, and the distance metrics, RMSD and GDTTS that relate the current snapshot to the starting model at time zero. For each snapshot the model is asked to predict whether an improved conformational state, no-change in conformational state or a decreased conformational state with respect to the reference crystal structure is sampled. The output softmax vector, and the ground truth assignment, are used to compute the weighted cross-entropy loss function. The model is trained with the Adam optimizer [39]. More details of the training procedure can be found in the SI text.

## Data and source code availability

The source code of the model as well as an example of how to train it is available from https://github.com/OneAngstrom/DeepTrajectory. The data used in this work is available for download from https://zenodo.org/record/1183354. This data contains the PDB files of the raw trajectories, starting models and reference PDB structures; and a comma separated file that contains the pre-computed trajectory features and labels.

## Acknowledgements

We would like to thank Robert Jenkins, Esther Wershof, David Jones and Cen Wan for the useful comments and discussions. This work was supported by the Francis Crick Institute which receives its core funding from Cancer Research UK (FC001003), the UK Medical Research Council (FC001003), and the Wellcome Trust (FC001003). We gratefully acknowledge the support of NVIDIA Corporation with the donation of the GPUs used for this research.

## Author information

### Author contributions

E.P. and P.A.B. designed the research and wrote the manuscript. E.P conducted the experiments and analysed the data.

## Supplemental Information

### Model Definition and Training

The model learns via a supervised learning-task to assign the class *y* from a set of given input features, *x*, for each time-point, [*τ*]_*i*_, of a trajectory *v*. Here, the three possible classes are improved, no-change and decreased. The ground truth assignment, denoted as *y′*, is then formalized such that

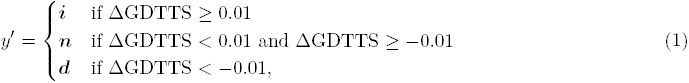

where *i, n* and *d* represent one-hot encoded probability vectors [1, 0, 0], [0, 1, 0] and [0, 0, 1], respectively. The variable ΔGDTTS is the difference between GDTTS from the starting model at time-point zero and the GDTTS from the snapshot at time-point *t* as computed to the reference crystal structure. A negative ΔGDTTS value reflects a decrease in model quality and a positive ΔGDTTS value an increase in model quality.

The model is based on an RNN with GRU that adaptively learns long and short term dependencies of inputs to assign the class *y* [19]. The layout of the RNN is illustrated in Fig. S1A, where starting from the input sequence *x*_0_, *…, x*_*t*_ the predictions *y*_0_, *…, y*_*t*_ are produced via stacking hidden layers of GRU cells and by a layer with a softmax activation function Ω that normalizes the output *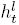*(at time *t* of the last layer *l*) to a probability vector *y* such that

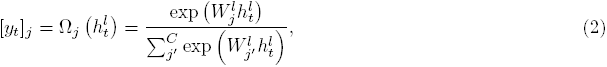

for all *j* = 1, *…, C* classes, where *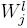* are the rows of the weight matrix of the last layer. The activation and its output, *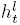* in layer *l* at time *t*, of a GRU cell is computed as

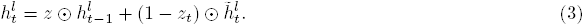

This represents a linear interpolation of the activation at time-point *t* − 1 denoted as *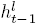* and its candidate activation 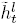. The update gate, *z*, controls how much the cell updates its state, such that

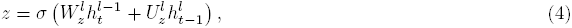

where the activation function *σ* is sigmoidal. The candidate state 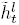 is computed such that

Here, *ϕ, r* and ⊙ denote a hyperbolic tangent activation function, a reset gate and an element wise multiplication, respectively. The reset gate, *r*, is computed with the same formulation as *z* but different weight matrices, i.e.

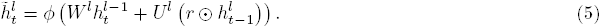

An illustration of these equations is shown in Fig. S1B.

During training the weight matrices *W*^*l*^, *U*^*l*^ *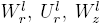* and *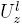* are learned for each layer *l*. The weight matrices are shared through time *t*. The loss function *L*

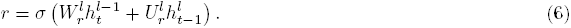

is minimized during training and represents the weighted cross entropy. The vector ***ω*** encodes the weights for classes *j* = 1, *…, C*. The objective of this classifier is to achieve a high precision for the improved class (i.e. reducing the false positive rate) and a high recall for the decreased class (i.e. reducing false negative rate). This is achieved by setting ***ω*** = [0.05, 1, 10].

The RNN model is trained with the Adam optimizer [39] on input features *x* from the set of trajectories *v* selected for training. In order to achieve one sequential input for the training, all *n* trajectories are concatenated to size *τ × n*. The training is performed on *k* mini-batches *b* of the data where in each training iteration the batch *b*_*k*_ continuous without overlap to the previous batch iteration till the epoch is finished. This process is visualized in Fig. S1C and S1D.

**Fig. S1:**
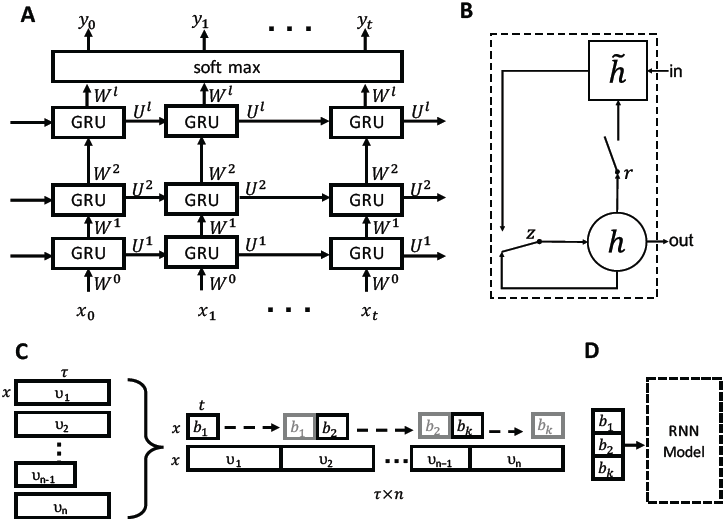
RNN model description. (A) Schematic overview of the RNN with GRU cells. (B) GRU cell, visualisation of Equations 3, 4, 6 and 5. (C) & (D) Visualisation of the trajectory data *v* and the process of mini-batch creation and propagation to the RNN.

## Data Set

The trajectory data originates from our own laboratory’s refinement method in CASP11 and CASP12 for which the reference crystal structure is available in the PDB. These targets are listed in SI Table S4 The detailed description of the sampling process is described in SI Section”Sampling procedure”. In total, the trajectory data consists of 3419 ns cumulated simulation time and 1, 709, 704 snapshots with Δ*t* = 2 ps from 30 CASP11 and 12 CASP12 targets for which crystal structures were available. A detailed overview of the generated trajectories for each target and their snapshot composition is provided in SI Table S5.

## Computation of Molecular Descriptors and Feature Construction

In total 19 features were used. 17 of these features originate from ten different potential energy functions and two features are the distance metrics GDTTS and RMSD that measure for each snapshots the difference to the starting model. All molecular descriptors are normalized per target to zero mean and unit standard deviation. The complete list of features is given in SI Table S1.

## Cross-Validation

The CV set is made up of 7 folds, where for each fold the training set consists of trajectories of 36 targets and the validation set for 6 targets. The assignment of a proteins trajectories to a fold is random. However, the relative distribution of classes of snapshots between training and validation set is enforced to be similar with a maximum difference of 6 percent as shown in Table S2 columns I, N and D. A detailed overview of each targets fold assignment is given in SI Table S5.

## Model Hyper-Parameter

The RNN model was trained for every fold of the CV for 300 epochs with the following hyper-parameter: sequence length = 30, batch-size = 50, internal size = 1024, number of layers = 3, learning rate = 0.0001 and dropout with p-keep = 0.9.

## Baseline Model

The RNN model was compared to the following baseline models:

**Random Forest [22]:** The training of the classifier uses 500 trees where samples are bootstrapped and the gini impurity criterion is used to judge the quality of the split when building the trees. No restriction for the maximum depth of the tree is imposed, however, for each internal node in a tree the minimum sample size must be greater than 30. The number of features for each tree is 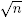 where n=19, i.e. the total number of features.

**K Nearest Neighbour [23]:** Number of neighbours and the leaf-size was set to 5 and 30, respectively. A uniform distribution where all points are weighted equally in each neighbourhood was chosen. The algorithm to search for the nearest neighbours was set to ‘auto’ where the best algorithm from ball-tree [46], kdtree [47] and a brute force approach was selected for fitting the model with the Minkowski distance metric with *p* = 2, which is equivalent to the Euclidean distance.

**Logistic Regression [24]:** Fitting of the model is performed with L2 regularization with a strength of 1.0 and with a tolerance of 1*e -* 4 as the stopping criteria. In order to make the multi-class predictions with LR, the training task is translated into a binary classification problem where for each label a fit of the LR is performed.

The python package scikit-learn [48] in version 0.18.1 was used to perform the training and testing. The same features, class-labels and CV folds as shown in Table S2 were used.

## Classifier Performance Metrics

In order to quantify the performance of the RNN and to compare it to other classifiers the metrics recall, precision and F1 (i.e. harmonic mean between precision and recall) were computed. For these three metrics the performance from all three classes are reported. These metrics are defined as follows:

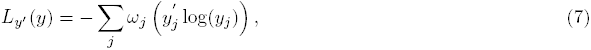

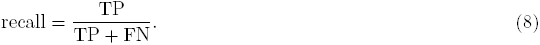

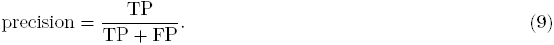

Where TP is the number of true positives, FN the number of false negatives and FP the number of false positives.

## Structural Model Assessment Metrics

The model quality of the snapshots is quantified by the two metrics GDTTS and C*α*-RMSD which are defined as follows

**RMSD:** The root mean square deviation quantifies the disagreement of the predicted model to the reference structure. A lower value indicates a better fit to the reference structure. The definition is such that

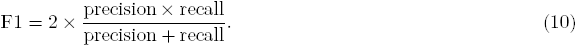

where ***v***, ***w*** are the set of atom coordinates for the model and reference structure, respectively. An optimal superimposition of ***v*** to ***w*** is performed prior to RMSD calculation.

**GDTTS:** The global distance test total score is a model quality metric that evaluates quality based on the percentage of residues under different distance cutoffs given by

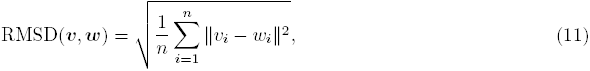

where *P*_*n*_ is the percentage of residues below distance cutoff *n* in Å with respect to a reference structure. A higher value indicates a better fit to the reference structure.

## Sampling procedure

The targets and starting models for the refinement simulations are given by the CASP committee and originate from rounds 11 and 12. These structures represent a diverse set of proteins with different folds and initial model quality ranging from 30 to 90 GDTTS.

The trajectories used for training and testing the RNN originate from 5 different MD-based sampling procedures. These are known as (i) Unrestrained sampling (no rst), no position or distance restraints to the residues of the starting are applied; (ii) position restraint sampling (point rst), position restraints to the C*α*-atom of structurally conserved residues are applied; (iii) distance restraint sampling (dist rst), residue-residue distance restraints are applied to residues that are structurally conserved; (iv) metadynamic sampling with exclusive residue-residue contacts (cm excl), sampling with position restraints and in contact map space (CMS) with metadynamics of unique residue-residue contacts; (v) metadynamic sampling with minimum distance residueresidue contact (cm min), sampling with position restraints and in CMS of residue contact pairs of the minimum initial distance.

## Position and distance restraint generation and sampling

The generation of restraints is outlined in Fig. S2A. For every CASP refinement target models from all participating predictors were downloaded. The number of models varies from target to target, but on average 180 submissions were available. Often, a substantial part of these submissions were physically implausible, i.e. they contained long extended stretches. In order to avoid including these into the analysis a 10 Å C*α*-RMSD cutoff to the provided starting model was applied.

From this filtered set, position and distance restraints for structurally conserved regions were generated and applied to C*α*-atoms. Position restraints were applied if the per-residue C*α*-RMSF calculated from the filtered set is *<* 3 Å (see Fig. S2C for an example). In order to determine conserved residue-residue distances all possible combinations of C*α*-C*α* pairs were measured and distance restraints were applied if all of the following criteria are true: a) the C*α*-C*α* pairs are at least 5 residues apart; b) the C*α*-C*α* distance is below 9 Å; c) the standard deviation of the distance is below 1 Å (see Fig. S2D for an example).

For each target three different simulation setups are executed: a) 3 ns long MD run without restraints, replicated 8 times; b) 3 ns long MD run with position restraints, replicated 8 times; c) 3 ns long MD run with distance restraints, replicated 8 times (see Fig. S2B). All MD simulations were computed with GROMACS, using version 4.6 [49], and the G54a7 force field [50]. For all initial target structures hydrogen atoms were added and the systems were neutralized with Na+ and Cl*-* counter ions. A cubic simulation system with a 12 Å buffer between the edge of the box and the protein was solvated with TIP3P water molecules [51]. All targets were then subject to an energy minimisation using the steepest decent algorithm with a maximum of 50000 steps. This was followed by an equilibrium phase to relax the structure and its solvent. MD simulations (a) and (b) were subject to a 2 step equilibrium protocol where all heavy atoms were position restrained by a force of 1000 kJ mol*-*1nm*-*1 throughout the equilibration. In the first phase an NVT equilibration of the system was performed to increase the temperature from 0 K to 300 K in 100 ps using V-rescale [52] for temperature coupling.

**Fig. S2:**
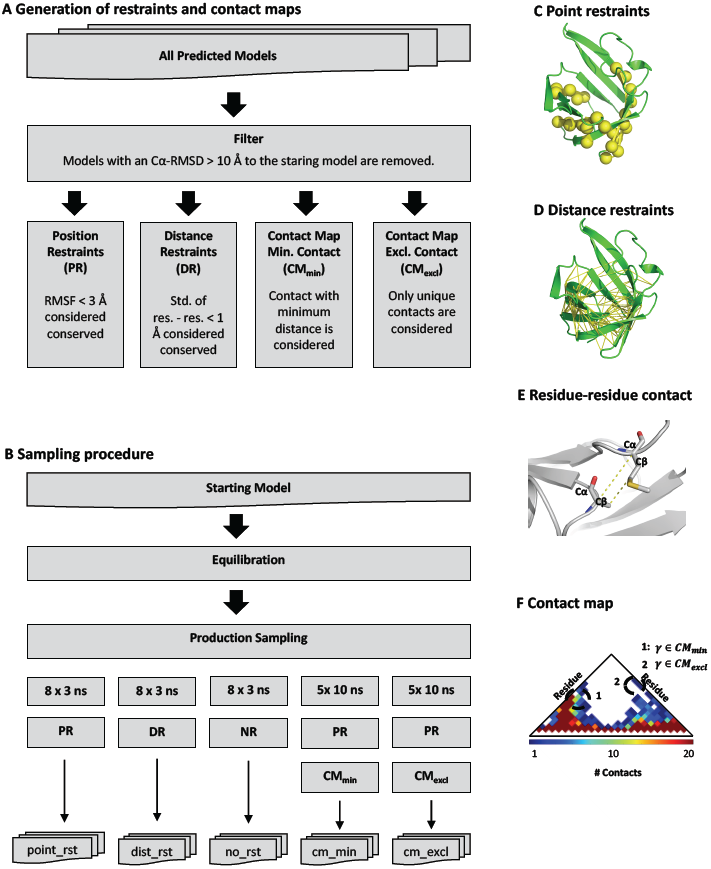
Sampling Protocol. (A) Flowchart outlining the generation of the restraints and contact maps. (B) Flowchart outlining the different MD sampling procedures (C) Example of point restraints applied to a protein. (D) Example of residue-residue distance restraints applied to a protein. (E) Definition of residue-residue contact (F) Contact map definition for CM_exl_ and CM_min_.

**A Generation of restraints and contact maps**

The second phase consisted of a 300 ps long NPT equilibration of the system’s pressure to 1 bar using Parrinello Rahman pressure coupling [53]. For MD simulation (c) a second NPT equilibration was applied, where the first step consisted of a 200 ps long equilibration with full heavy atoms position restraints and distance restraints, and the second step of a 200 ps equilibration with distance restraints only.

For all simulation setups a leap-frog integrator with a Δt of 2 fs was used and coordinates, velocities, energies, and forces were saved every 2 ps. Long range electrostatic interactions were treated with the Particle Mesh Ewald method [54] with a cutoff of 10 A. Temperature and pressure coupling were controlled by the V-rescale and the Parrinello-Rahman method and were set to 300 K and 1 bar, respectively.

### Contact map generation and sampling

All available models from participating predictors of a target are downloaded from the prediction center server. Each model is compared to the starting model and the C*α*-RMSD is calculated, models with a C*α*-RMSD *>* 10 A are removed from the set. This was done in order to remove outliers from the set.

The filtered set is used to determine structurally conserved residues. These are identified by computing the per residue root mean square fluctuation (RMSF) of C*α* atoms. Residues with a RMSF *<* 3 Å are considered conserved and movements are restraint during the sampling process.

From the structures in the filtered set of CASP predictions, residue-residue contacts are identified with a C*α* or C*β* distance below 8 A (Fig. S2E), with the exception of direct neighbours, which are removed from the list. From these contacts two contact maps (CM) are generated, namely CM_exl_ and CM_min_ (Fig. S2F). CM_exl_ contains contacts that are exclusive to one model from the filtered set, whereas the map CM_min_ contains contacts with the lowest C*α*/C*β* distance. From these CMs we can define two CVs describing the CMS:

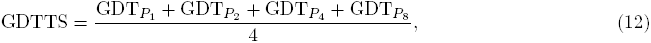

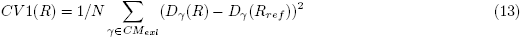

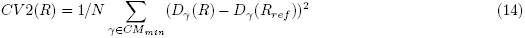

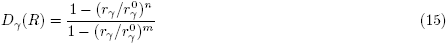

The sigmoid distance function *D*_*γ*_ (*R*) is used to quantify the formation of a contact *γ* in structure *R*, where *r*_*γ*_ is the contact distance in structure *R* and *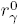* is the contact distance in reference structure *R*_ref_ which denotes to one of the models from the filtered set of CASP12 models where the contact was observed. Variables *n* and *m* are constant and set to *n* = 6 and *m* = 10.

The preparation of the starting model prior to the sampling process follows a GROMACS standard procedure where the system is solvated, energy minimized and equilibrated for 300 ps. The sampling with metadynamics in CMS is performed at 300 K for 10 ns with 5 replicas for each CM definition, resulting in 100 ns sampling data for each target. The sampling of the CMS was performed with the GROMACS plug-in PLUMED2 [55] where a Gaussian addition is deposited every 2 ps with *σ* = 0.5, a bias factor of 10 and an initial height of 5 kJ*/*mol.

---

**Fig. S3:**
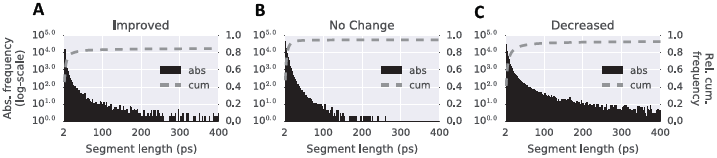
Histogram of the continues segmentation length of the three different states in all trajectories. Frequency as a function of continuous segment length with (A) improved quality, (B) no change in quality and (C) decreased quality.

**Fig. S4:**
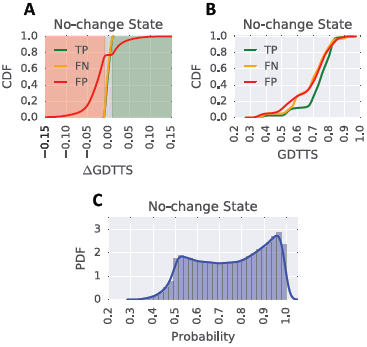
(A) Cumulative distribution of true positive (TP), false negative (FN) and false positive (FP) predictions for the no-change state as a function of ΔGDTTS. The background colours red, gray and green indicate the ΔGDTTS regions for the improved, no-change and decreased states, respectively. (B) Cumulative distribution of true positive (TP), false negative (FN) and false positive (FP) predictions for the no-change state as a function of GDTTS. (C) Show the distribution of assigned probabilities for no-change state predictions.

**Fig. S5:**
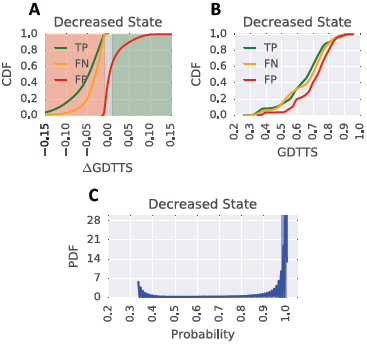
(A) Cumulative distribution of true positive (TP), false negative (FN) and false positive (FP) predictions for the decreased state as a function of ΔGDTTS. The background colours red, gray and green indicate the ΔGDTTS regions for the improved, no-change and decreased states, respectively. (B) Cumulative distribution of true positive (TP), false negative (FN) and false positive (FP) predictions for the decreased state as a function of GDTTS. (C) Show the distribution of assigned probabilities for decreased state predictions.

**Table S1:** RNN features. Table lists the feature name, the descriptor that produced the feature and the reference for the descriptor.

| Feature | Descriptor | Reference |
| --- | --- | --- |
| N_DDFIRESUM | DFIRE | [40] |
| N_DDFIRETERM1 | DFIRE | [40] |
| N_DDFIRETERM2 | DFIRE | [40] |
| N_DDFIRETERM3 | DFIRE | [40] |
| N_DDFIRETERM4 | DFIRE | [40] |
| N_DOPE | DOPE | [41] |
| N_DOPE_HR | DOPE | [41] |
| N_RMSD_SM | RMSD to starting model | NA |
| N_GDTTS_SM | GDTTS to starting model | NA |
| N_DOOP | DOOP | [42] |
| N_CALRW | calRW | [43] |
| N_CALRWP | calRWplus | [43] |
| N_GOAP | GOAP | [44] |
| N_GOAPAG | GOAP | [44] |
| N_BOND | Modeller | [45] |
| N_ANGLE | Modeller | [45] |
| N_DIHEDRAL | Modeller | [45] |
| N_IMPROPER | Modeller | [45] |
| N_MOLPDF | Mol. PDF | [45] |

**Table S2:** Cross validation summary. Summary of each fold of the 7-fold CV of the trajectory data. Shown are the number of targets (# TR), number of trajectories (# Trj), number of snapshots (# Snap.), percentage of snapshots with improved quality (I (%)), percentage of snapshots with no change in quality (N (%)), percentage of snapshots with decreased quality (D (%)).

| Training Set |  |  |  |  |  |  |
| --- | --- | --- | --- | --- | --- | --- |
| Fold | # TR | # Trj. | # Snap. | I (%) | N (%) | D (%) |
| 0 | 36 | 780 | 1,463,580 | 8.38 | 15.00 | 76.62 |
| 1 | 36 | 772 | 1,451,572 | 8.39 | 14.07 | 77.54 |
| 2 | 36 | 772 | 1,451,572 | 8.54 | 14.00 | 77.46 |
| 3 | 36 | 772 | 1,451,572 | 7.47 | 14.38 | 78.14 |
| 4 | 36 | 780 | 1,462,780 | 8.09 | 14.46 | 77.45 |
| 5 | 36 | 782 | 1,499,782 | 8.22 | 14.90 | 76.88 |
| 6 | 36 | 766 | 1,477,366 | 8.35 | 14.87 | 76.78 |

| Validation Set |  |  |  |  |  |  |
| --- | --- | --- | --- | --- | --- | --- |
| Fold | # TR | # Trj. | # Snap. | I (%) | N (%) | D (%) |
| 0 | 6 | 124 | 246,124 | 7.16 | 11.74 | 81.10 |
| 1 | 6 | 132 | 258,132 | 7.17 | 17.14 | 75.70 |
| 2 | 6 | 132 | 258,132 | 6.34 | 17.49 | 76.17 |
| 3 | 6 | 132 | 258,132 | 12.34 | 15.35 | 72.31 |
| 4 | 6 | 124 | 246,924 | 8.90 | 14.93 | 76.17 |
| 5 | 6 | 122 | 209,922 | 8.12 | 11.87 | 80.01 |
| 6 | 6 | 138 | 232,338 | 7.31 | 12.36 | 80.33 |

**Table S3:**
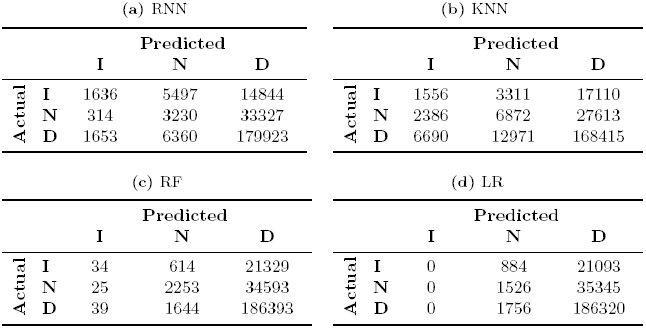
Confusion matrix. The four different sub-tables show CV (validation-fold 4) of the predicted and actual class assignment for improved (I), no change (N) and decreased (D) for (A) RNN, (B) KNN, (C) RF and (D) LR.

**Table S4:**
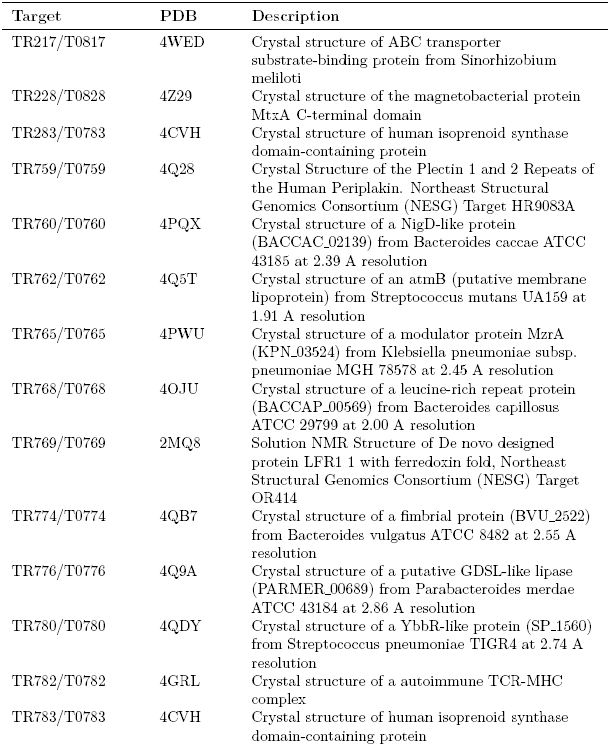
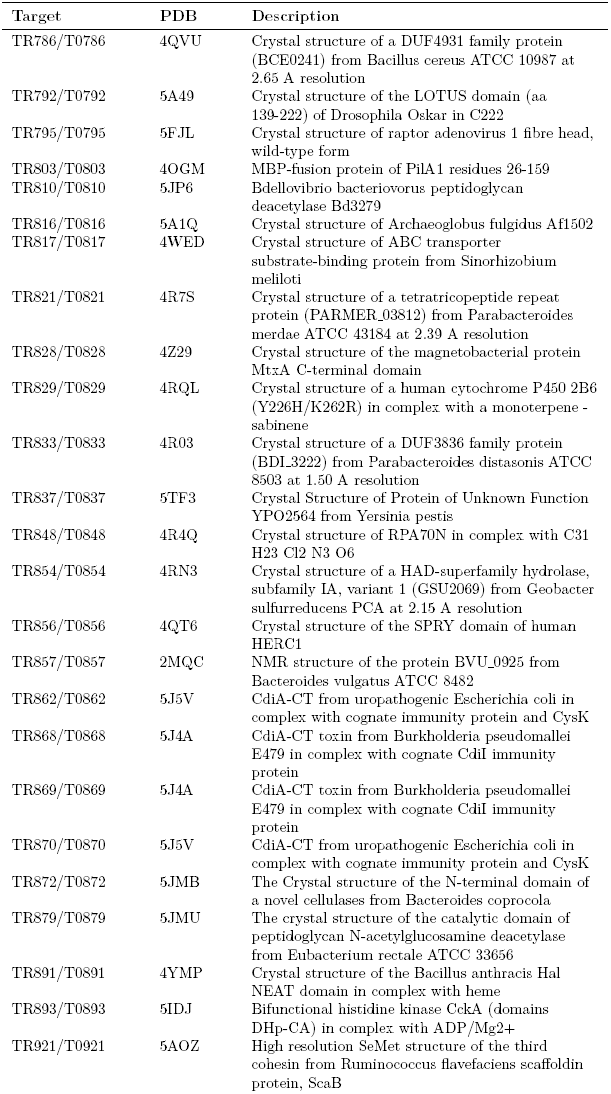
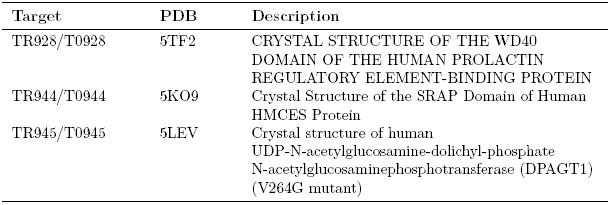
Protein target overview of which several MD trajectories were generated

**Table S5:**
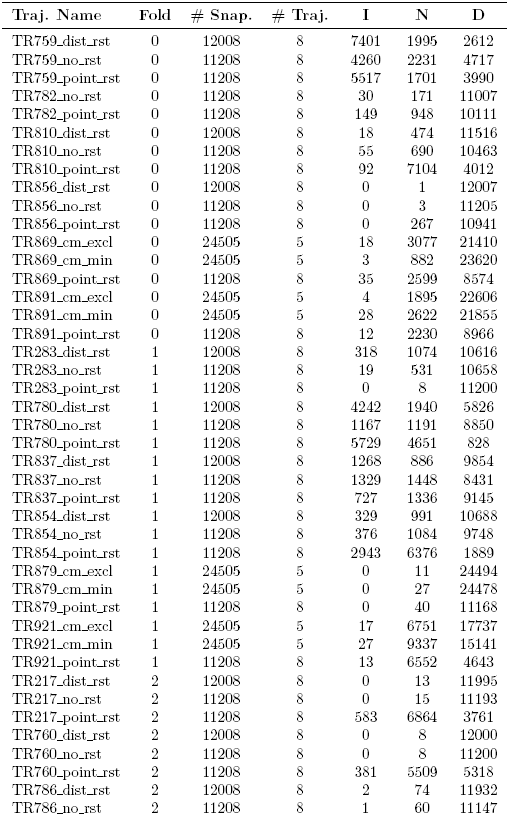
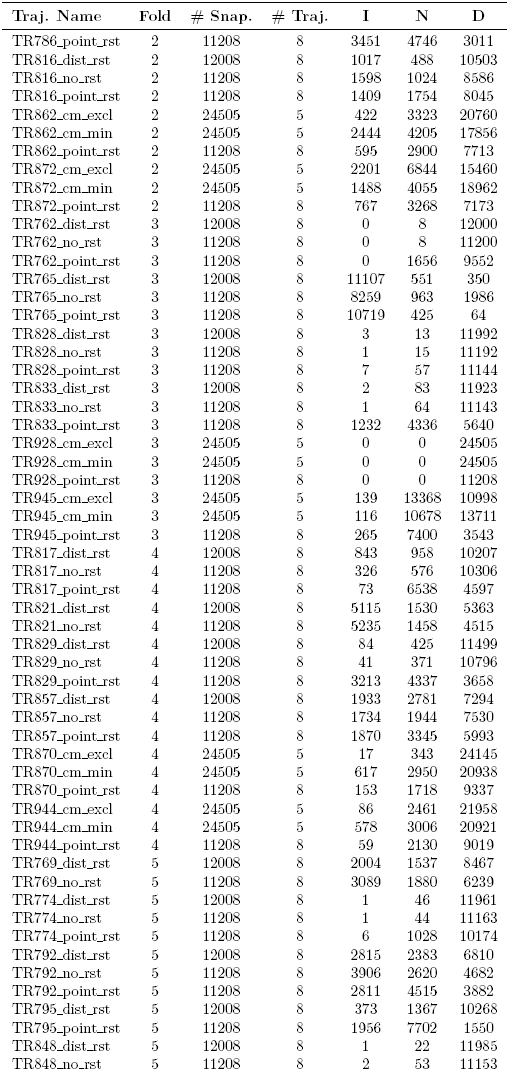
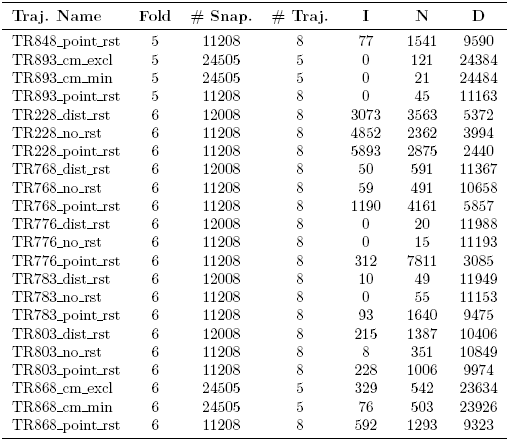
Trajectory and cross validation overview for all protein targets.

